# Early nascent polypeptide dynamics are coupled to the flexibility of the ribosomal tunnel constriction

**DOI:** 10.64898/2026.03.10.710814

**Authors:** Hugo McGrath, Michaela Černeková, Michal Kolář

**Affiliations:** Department of Physical Chemistry, University of Chemistry and Technology, Technická 5, Prague, Czech Republic

**Author notes:** equal contribution (Hugo McGrath, Michaela Černeková).

## Abstract

The ribosomal exit tunnel, through which all nascent polypeptides emerge, is formed primarily by ribosomal RNA. Still, ribosomal proteins contribute roughly one quarter of the tunnel walls. In particular, proteins uL4 and uL22 define the narrowest region of the tunnel. This constriction, enriched in basic residues, mediates the earliest protein–protein contacts experienced by a nascent polypeptide. Here, we characterize its conformational dynamics by analyzing 222 *Escherichia coli* ribosome structures from the Protein Data Bank and by performing unbiased all-atom molecular dynamics simulations of the complete bacterial ribosome with nascent polypeptides of varying length and composition. In simulations of the empty ribosome, the constriction can transiently narrow below the diameter of a water molecule and thus fully occlude the tunnel. At other times it opens wide enough to accommodate a narrow *α*-helix. The presence of even a short nascent polypeptide shifts the constriction toward wider conformations by approximately 0.2 nm, revealing an adaptive response of the tunnel to its contents. Across all simulated systems, the N-terminal formylmethionine preferentially associates with the tunnel wall at the uL22 side. Our findings replace qualitative description of the tunnel dynamics with a quantitative one. The constriction site functions as a flexible gate, with implications for nascent polypeptide translocation, action of macrolide antibiotics, and the translational ramp.

## 1 Introduction

During translation, all proteins emerge from the ribosome through a tunnel in the large ribosomal subunit. The tunnel accommodates a few dozen residues of the nascent polypeptide (NP) before it is long enough to reach the ribosome surface, where it begins interacting with the rest of the cell.^1,2^ During these early moments, the NP interacts with its environment, and through these interactions it contributes to translational regulation.^3^ In addition to interactions with the tunnel walls, the NP may be exposed to metabolites and other small molecules, such as antibiotics, present in the tunnel.^4,5^ Nevertheless, the primary environment of the nascent polypeptide is aqueous.

The ribosomal tunnel spans approximately 10 nm from the peptidyl transferase center (PTC), where peptide bond synthesis takes place, to the ribosome surface. The tunnel walls are composed predominantly of ribosomal RNA (rRNA), with several distinct exceptions represented by ribosomal proteins (r-proteins), among which uL4, uL22, uL23, and uL24 are the most conserved. In *E. coli*, r-proteins account for roughly 27.5% of the tunnel wall surface although the rRNA-to-r-protein ratio varies along the path. Notably, the PTC is formed exclusively by rRNA. ^6^

The tunnel exhibits distinct electrostatic properties that facilitate NP translocation. ^7,8^ Its cross-sectional area varies considerably along the path. One major constriction has been identified in the tunnels of bacterial ribosomes, while two constrictions occur in those of eukaryotic ribosomes.^9^ Based on structural models derived by X-ray crystallography and cryogenic electron microscopy (cryo-EM), the constriction is approximately 0.8 nm wide – a property we examine in greater detail in this work.

The main constriction site (CS), present in ribosomes across all three domains of life, is formed by the extended loops of r-proteins uL4 and uL22. The globular domains of these proteins are located near the ribosome surface and represent evolutionarily younger parts of the proteins.^10^ The evolutionarily older extended loops protrude deep into the large ribosomal subunit (LSU), reaching the tunnel interior. The distance between the PTC and the CS is approximately 2.6 nm (the first third of the exit tunnel). This early part of the tunnel accommodates short NPs before they traverse the CS. The NP path from the PTC to the CS is lined exclusively by rRNA, CS is therefore the site of the first protein–protein interactions in the life of a newly synthesized protein. The precise circumstances of this initial NP–CS contact remain unknown.

The amino-acid composition of these extended loops is enriched in basic residues, many of which are conserved across species, suggesting an evolutionarily important function. Among the most conserved residues is Gly91 of uL22, which likely contributes to the high conformational flexibility of the extended loop. Several mutations in uL22 have been identified that substantially affect bacterial fitness.^11^ This region of the tunnel also harbors binding sites for clinically important classes of antibiotics. Macrolide antibiotics bind in the vicinity of the uL22 extended loop, where they can modulate its conformation and discriminate among NP sequences. ^12,13^ Mutations in both uL4 and uL22 have also been linked to macrolide resistance. Two distinct resistance mechanisms have emerged from previous studies: mutations that reduce macrolide binding affinity, predominantly in uL4 but also in uL22,^14,15^ and mutations that widen the tunnel to allow NP bypass even while the drug remains bound. ^13,16^ In this work, we investigate the flexibility of the CS in bacterial ribosomes and the factors that govern it. Our primary aim is to move beyond the single-conformation structural models derived from experiment and the qualitative picture that has reduced to a rather intuitive statement that the CS is flexible. We acknowledge that the static tunnel assumption, which has been widely used for example to study cotranslational folding ^17–20^ and protein escape from the tunnel^8,21–25^ under various circumstances, intentionally simplified the problems under investigation. However, the extent to which this assumption holds has remained unclear. Here, we perform a thorough analysis of an ensemble of available experimental structural models to quantify the conformational variability of the CS. We complement this analysis with atomistic molecular dynamics (MD) simulations, which have become a powerful tool for studying ribosomes, as recently reviewed. ^26,27^ Using probably the most accurate computational model of the entire atomistic fully dynamic ribosome in an explicit aqueous environment, we characterize the sub-microsecond dynamics of the CS and demonstrate that how flexible it actually is.

## 2 Methods

### 2.1 Protein Data Bank set

Cryo-EM and X-ray crystallography structures of either whole *E. coli* ribosomes or at least their LSU with resolution below 4.0 Å were downloaded from the Protein Data Bank. Entries with mutated uL4 or uL22 were omitted. Structures with incomplete tips were also omitted. The resulting data set contained 222 structures with r-protein uL4 and uL22 in various chemical contexts (nascent polypeptides, antibiotics, metabolites, etc.). Their PDB codes are available in the accompanying Data repository. R-proteins uL4 and uL22 were extracted from the structural models and analyzed. After superposition of the C_*α*_ atoms of both r-proteins, performed in Pymol, ^28^ we calculated the root-mean-square fluctuation (RMSF) of two sets of atoms, as a proxy of their conformational flexibility. First, only the C_*α*_ atoms of the tips were considered. Next, we calculated the RMSF of the terminal non-hydrogen atoms of the side chains of each residue.

### 2.2 Simulated systems

The simulated model of the *E. coli* ribosome was taken from our previous work.^29^ Briefly, the model is based on a cryo-EM structure reported by Fischer *et al*. ^30^ It contains an A-site tRNA^Phe^ and a P-site tRNA bearing a nascent polypeptide. Five different systems were simulated, differing in the NP, as depicted in Fig. 1. The shortest system, denoted Empty, contained only fMet on the P-site tRNA. Additional systems included poly-glycine or poly-alanine chains of either 5 or 11 residues, denoted Gly5, Gly11, Ala5, and Ala11. In all cases, the NP was terminated by an N-terminal fMet residue. An overview of the simulated systems is provided in Table 1.

**Table 1:** Summary of the simulated ribosome systems. Each system contained an A-site tRNA^Phe^, P-site tRNA with the peptide.

| name | peptide | replicas | ns/replica |
| --- | --- | --- | --- |
| Empty | fMet | 8 | 250 |
| Gly5 | (Gly)<sub>5</sub>-fMet | 8 | 250 |
| Gly11 | (Gly)<sub>11</sub>-fMet | 8 | 250 |
| Ala5 | (Ala)<sub>5</sub>-fMet | 8 | 250 |
| Ala11 | (Ala)<sub>11</sub>-fMet | 8 | 250 |

**Table 2:** Summary of the experimental structures with the narrowest CS as annotated in Fig. 3. Constriction-site width (CSw) and center-of-geometry distance (CoGd) are in nm, resolution in Å.

| PDB code | CSw [nm] | CoGd [nm] | res. [Å] | year | content |
| --- | --- | --- | --- | --- | --- |
| 4V5B | 0.39 | 1.78 | 3.74 | 2014 | PDF helix bound to 70S ribosome<sup>53</sup> |
| 5KPX | 0.46 | 1.54 | 3.90 | 2016 | RelA bound to 70S ribosome<sup>54</sup> |
| 5KPW | 0.86 | 1.56 | 3.90 | 2016 | RelA bound to 70S ribosome<sup>54</sup> |
| 6GBZ | 0.72 | 1.49 | 3.80 | 2018 | LSU assembly intermediate<sup>55</sup> |
| 6O9J | 0.59 | 1.69 | 3.90 | 2019 | 70S ribosome<sup>56</sup> |
| 6WNT | 0.50 | 1.50 | 3.10 | 2020 | 70S ribosome lacking 5S rRNA<sup>57</sup> |
| 6WNV | 0.57 | 1.69 | 3.50 | 2020 | LSU lacking 5S rRNA<sup>57</sup> |

**Figure 1:**
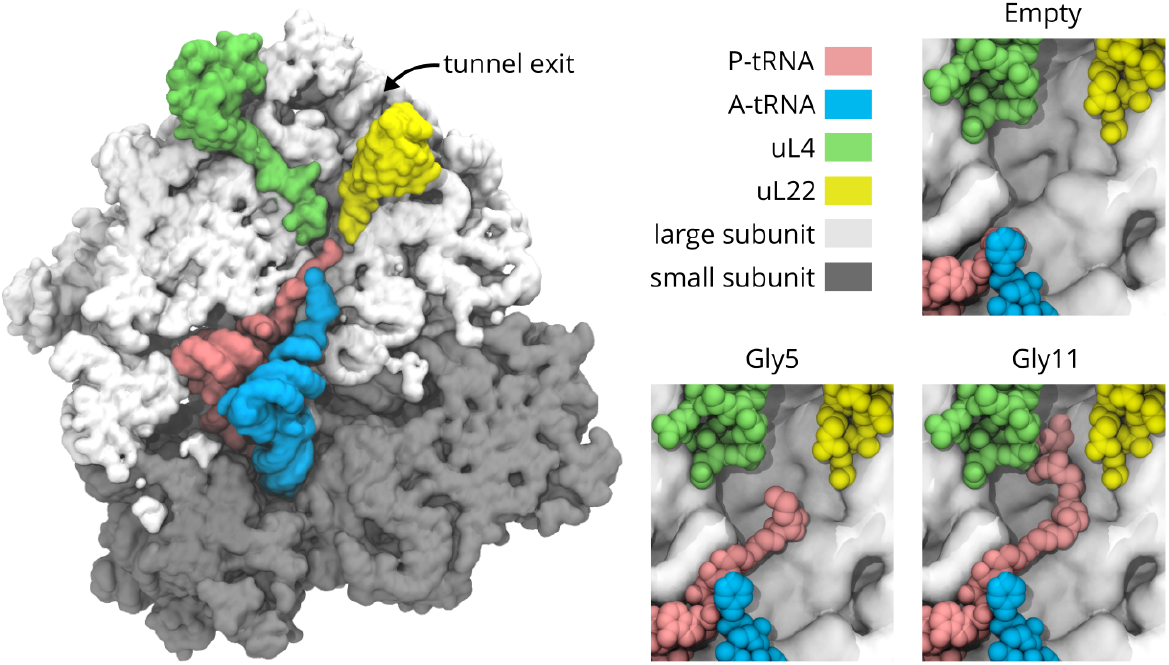
Simulated systems. Cross-sectional view of the *E. coli* ribosome, with the small and large subunits shown in dark and light gray, respectively. The A-site tRNA is shown in blue, the P-site tRNA in pink, and the r-proteins uL4 and uL22 in green and yellow, respectively. The right panels show detailed views of the PTC with the initial conformations of nascent polypeptides of three different lengths.

The initial peptide conformations were taken from a cryo-EM model of the ribosome stalled by the TnaC peptide (PDB 4V5H^31^). The TnaC sequence was converted to polyglycine or poly-alanine in PyMOL and truncated as required. Note that TnaC causes translational arrest only in the presence of free tryptophan residues.^31^ Thus, the absence of Trp, truncation of the peptide, and conversion to a homorepeat sequence ensure that the NP does not retain the stalling capability of TnaC. An fMet residue was manually added to the N terminus of each NP. The NP was positioned within the ribosomal exit tunnel by superposition of the phosphorus atoms of the P-site tRNAs from structures 5AFI and 4V5H.

Each ribosome was put in a rhombic dodecahedral box with water so that the solute was a minimal distance of 0.85 nm away from the box faces. Periodic boundary conditions were applied in all three box dimensions. Structural Mg^2+^ ions were taken from the 5AFI cryo-EM model. Appropriate number of bulk water molecules was randomly replaced with K^+^ ions to reach electroneutrality. Excess KCl and MgCl_2_ were added to achieve a concentration of 100 mmol dm^*−*3^ and 10 mmol dm^*−*3^ respectively.

A combination of Amber family of force fields was used, namely ff12SB^32^ for proteins, ff99 with bsc0 and *χ*_*OL*3_ corrections for canonical RNA nucleotides^33–35^ and Aduri *et al*. parameters for non-canonical nucleotides^36^ (a few dozen in the entire system). SPC/E was chosen as the water model^37^ along with Joung and Cheatham ion parameters. ^38^ This combination of force-field parameters has been shown to perform well in previous simulation studies of the bacterial ribosome from our group^29,39^ and from others. ^40,41^ Alternate choices could also be possible, however.

### 2.3 Molecular dynamics simulations

All simulations were performed using the GROMACS 2024 simulation package.^42^ Prior to the production runs, the systems were carefully prepared through a series of equilibration simulations to avoid large conformational changes arising from spurious close interatomic contacts that may occur during system setup.

First, the solvent was energy-minimized using the steepest-descent algorithm for 50,000 steps, while solute atoms were restrained with harmonic position restraints (force constant 1000 kJ mol^*−*1^ nm^*−*2^). The solvent was then heated to 300 K over 50 ps using a 1 fs time step and the v-rescale thermostat.^43^ During this stage, the solute was position-restrained and maintained at 10 K. Initial velocities for all particles were drawn from the Maxwell– Boltzmann distribution at 10 K, and the temperature coupling constant *τ* was set to 0.5 ps. Next, pressure equilibration was carried out for 50 ps using the Berendsen barostat, ^44^ targeting 1 bar with a pressure coupling constant *τ*_*p*_ = 1 ps. Subsequently, in a 1 ns simulation, the temperature of the entire system (solute and solvent) was equilibrated to 300 K using the v-rescale thermostat (*τ* = 1 ps), while the pressure was equilibrated to 1 bar using the c-rescaling barostat^45^ with *τ*_*p*_ = 1 ps.

Position restraints were then gradually released over a 5 ns simulation at a reference temperature of 310 K. The release was controlled via an alchemical parameter (*λ*) applied to the restraining potential. In a subsequent 1 ns simulation, the time step was increased to 2 fs, and both the temperature and pressure coupling constants were set to 2 ps. This was followed by a 15 ns simulation in which the time step was increased to 3 fs by employing hydrogen mass repartitioning (HMR) with a factor of 2.^46^ Finally, during the production stage, an HMR factor of 3 was used, allowing the time step to be extended to 3.5 fs.

A total of eight independent production trajectories were generated at 310 K and 1 bar, differing only in the initial velocity assignments during the heating stage. Each trajectory was 200 ns long, and coordinates were saved every 10,000 steps.

In all simulations, Newton’s equations of motion were integrated using the leapfrog integrator. Bond lengths involving hydrogen atoms were constrained using the P-LINCS algorithm^47^ of order 6. Long-range electrostatics were treated with the particle–mesh Ewald method,^48^ using a real-space cutoff of 1 nm, an interpolation order of 4, and a grid spacing of 0.12 nm. Lennard-Jones interactions were truncated at a cutoff distance of 1 nm. The input data as well as a complete set of MD parameters are provided in the accompanying Data repository.

### 2.4 Analyses

The analyses were performed with custom Python (v3.11.5) scripts employing NumPy ^49^ (v1.24.3) and MDAnalysis^50^ (v2.6.1) with Matplotlib ^51^ (v3.7.2) for plotting and VMD^52^ (v1.9.2) for molecular visualization. The scripts are available online in an accompanying Data repository.

We analyzed eight independent trajectories for each of the five systems (Empty, Gly5, Gly11, Ala5, and Ala11). For simplicity, only non-hydrogen atoms were considered in the analyses. If not stated otherwise, the first third of the trajectories were left for systems relaxation and omitted, giving the latter two thirds for the analyses.

Before the analyses, the trajectories were iteratively aligned to the most rigid P atoms of the entire ribosome. The alignment procedure was as follows: initially, all P atoms were used for alignment. The RMSF values of these atoms were then calculated, and the 90% most rigid atoms were selected for the next iteration. This process was repeated five times, resulting in an alignment less influenced by the mobile, mostly peripheral regions of the ribosome.

Two metrics were defined to characterize the CS. The *CS width* is the distance between the closest non-hydrogen atoms in the uL4 and uL22 tips, typically involving atoms from the side chains of the two r-proteins. The *CoG distance* was defined as the distance between the centroids (*i. e*., centers of geometry, CoG) of the tip C_*α*_ atoms. For the purpose of both metrics, the tips were defined as residues 63–67 and 88–93 for uL4 and uL22, respectively. This definition was the same for the PDB ensemble and for the MD trajectories.

The Gibbs energies in the 2D space of CS width and CoG distance were obtained from probability density functions (PDFs). First, we estimated the PDFs as 2D histograms *H* normalized to unit area for each individual trajectory. Using bootstrapping with replacement, we resampled the set of 8 histograms 10,000 times. For each bootstrap replicate, the mean histogram was computed and converted to Gibbs energy as *G* = *−RT* ln *H*, and the profile was shifted so that the global minimum was set to 0 kJ/mol. The reported mean Gibbs energy profile and standard error (SE) were obtained as the mean and standard deviation, respectively, over the 10,000 bootstrap replicates. For plotting purposes, infinite values arising from regions of the 2D space not visited during the MD simulations were replaced by the maximum finite Gibbs energy value.

By integrating the PDF estimates, we obtained the percentage of frames where the CS width is below a given threshold. We calculated an average value as well as standard errors of the mean across 8 trajectories.

The atoms closest to each other in the CS in each frame were tracked and we report, for each residue, the percentage of frames in which it contributed the atom closest to the opposing protein. This was done for the aggregate of all trajectories for each system and both CS proteins.

End-to-end (E2E) distance calculations were performed for the NP conformations as the Euclidean distance of the C_*α*_ atoms of the terminal residues. The PDFs were obtained for each trajectory as histograms normalized to unit area.

The PDFs of non-hydrogen atoms’ positions of the peptide, uL4 and uL22 tips as well as water were calculated in a cylinder along the axis of the exit tunnel as shown in Fig. 2. The tunnel axis was defined by the centroid of residues 64 of uL4 and 92 of uL22 on one side and the centroid of the C-terminal NP residue on the other. The cylinder has a radius of 2 nm and height of 3 nm. The positions of the atoms in the cylinder are projected onto its base. For visualization purposes the projections were transformed to polar coordinates. The centroid of the residues 60–67 of uL4 was selected to represent 0^*°*^ of rotation around the cylinder axis. Complete trajectories were used for the projections.

**Figure 2:**
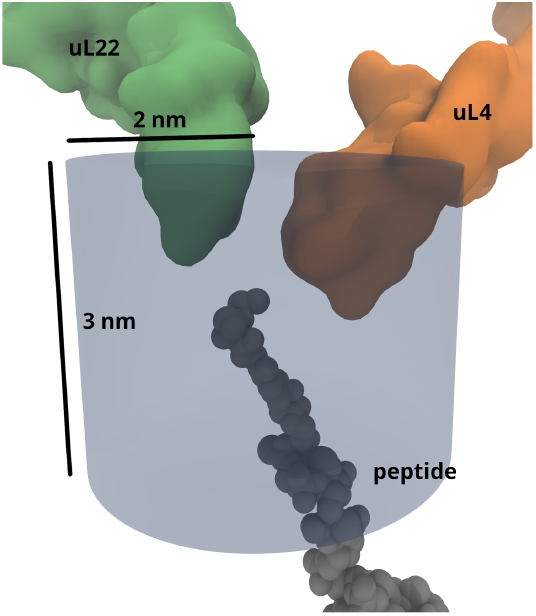
A cylinder with a radius of 2 nm and height of 3 nm oriented along the axis of the ribosomal tunnel used for water analysis.

## 3 Results and Discussion

### 3.1 Constriction site is flexible

We characterized the conformational dynamics of the CS using CS width and CoG distance. Both metrics highly fluctuate throughout the independent trajectories, as show for the Empty system in Fig. 3A. We constructed Gibbs free-energy landscapes as a function of CS width and CoG distance (Fig. 3B). All simulations were initiated from a peptide-free ribosome (PDB 5AFI ^30^) with A-tRNA-Phe and P-tRNA-NP occupying the PTC. In this structure, the CS width is 1.0 nm and the CoG distance 1.7 nm. Over 200 ns of unbiased MD simulation, the CS sampled a wide region of conformational space, underscoring its intrinsic flexibility. The global energy minimum of the Empty simulations closely matches the experimental values from 5AFI. Furthermore, the simulations span the full range of CS widths observed across 222 experimental *E. coli* ribosome structures (Fig. 3B, white points), supporting the validity of the simulation model. Because these experimental structures represent ribosomes in diverse biochemical contexts, we examined in detail those with the narrowest CS (Fig. 3B; Tab. 2). These structures share two notable features: comparatively low resolution (6WNT excepted) and substantial biochemical perturbations, such as the absence of 5S rRNA. This suggests that such perturbations may exert non-local conformational effects on the tunnel.

**Figure 3:**
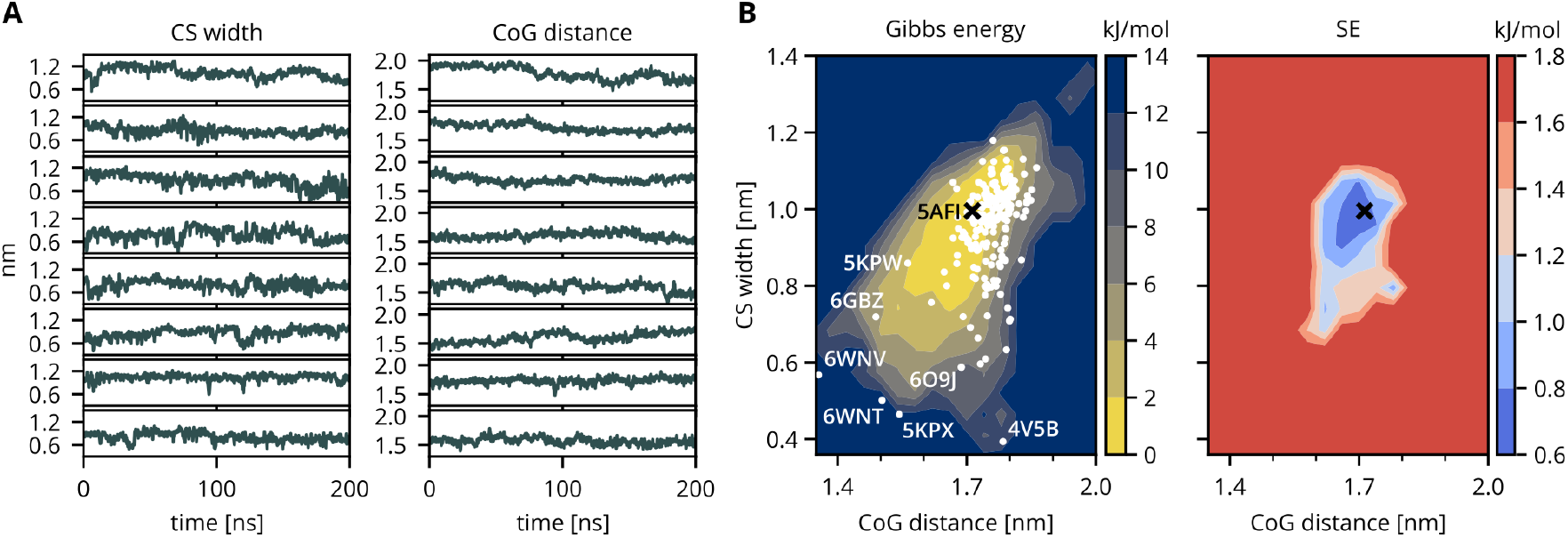
Constriction-site (CS) analysis of the Empty simulations. A) CS width and center-of-geometry (CoG) distance as a function of simulation time. B) Gibbs energy at 310 K as a function of CS width and CoG distance, color-coded from yellow to navy. The black cross marks the initial configuration derived from the cryo-EM structure of the ribosome with an empty tunnel (PDB 5AFI^30^). White points represent 222 experimental structures of *E*.,*coli* ribosomes. Structures with the narrowest CS are annotated with their PDB codes and summarized in Tab. 2. The standard error (SE) of the Gibbs energy is shown color-coded from blue to red.

To assess the per-residue contributions to CS flexibility, we calculated RMSF values for the tips of uL4 and uL22 across the 222 experimental structures (Fig. 4A). The residues exhibiting the greatest conformational variability within the exit tunnel are Trp60, Arg61, and Arg67 on the uL4 side, and Arg92 and Arg95 on the uL22 side. This analysis indicates that CS dynamics are dominated by side-chain rearrangements rather than backbone motion. The high flexibility of these charged and bulky residues suggests that electrostatic and steric interactions at the CS are potentially contributing to the gating of NP passage through the tunnel.

**Figure 4:**
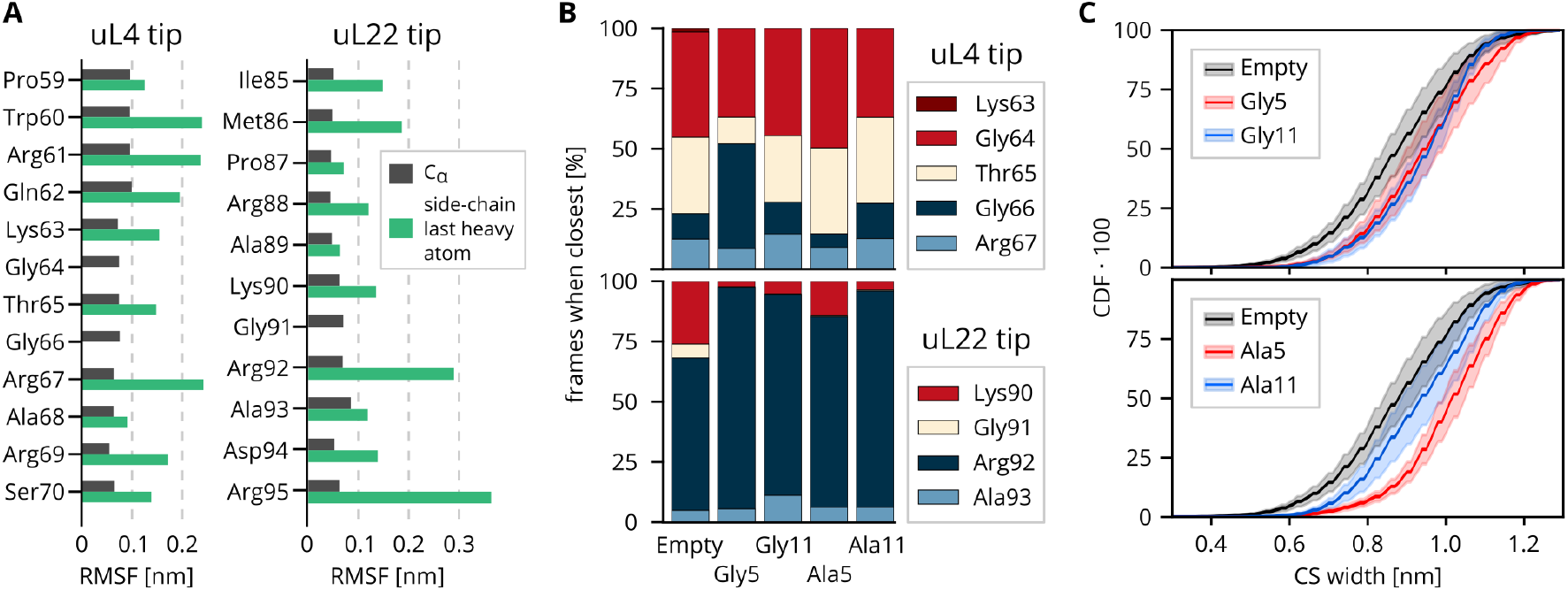
A) Root-mean-square fluctuations (RMSF) of C_*α*_ atoms (black) and the last non-hydrogen atoms of the side chains (green) of uL4 and uL22 tips calculated from the 222 experimental structure. B) Percentage of frames where a given residue of uL4 and uL22 is the closest to its counterpart. C) A cumulative distribution function (CDF) of the constriction-site (CS) width. The lines represent the mean values over the 8 independent trajectories, the shaded areas are standard errors of the mean.

We next identified which residue pairs form the closest contacts between uL4 and uL22 in the MD simulations (Fig. 4B). The narrowest points of the CS are most frequently defined by Gly64 and Thr65 on uL4, and Arg92 on uL22 – residues among the most conserved across prokaryotes and eukaryotes,^11^ suggesting that precise regulation of CS width is functionally important. Arg92 features prominently in both analyses: it is both highly flexible and the most frequent mediator of the closest uL4–uL22 contact. We therefore conclude that Arg92 plays a dominant role in CS width variation, sampling multiple side-chain conformations as it scans the uL4 tip.

### 3.2 CS may get closed

A natural question arising from the observed CS flexibility is whether the CS acts as a gate that opens and closes, potentially regulating the passage of the NP through the ribosomal tunnel. We addressed this by analyzing the Gibbs free-energy profiles of the Empty systems (Fig. 3) and of the systems with short NPs (Fig. 5). Furthermore, we calculated the cumulative distribution function (CDF) of CS width, which represents the fraction of simulation time spent below a given width threshold (Fig. 4C).

**Figure 5:**
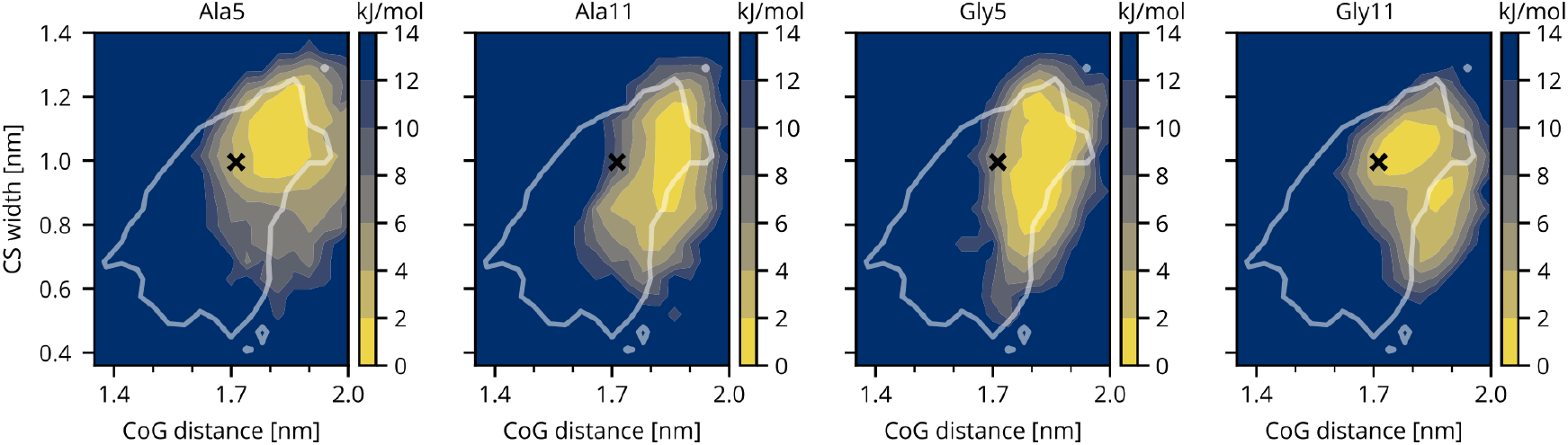
Gibbs energy profiles of systems with short nascent polypeptides at 310 K as a function of the CS width and the CoG distance. The black cross marks the initial configuration derived from the cryo-EM structure of the ribosome with an empty tunnel (PDB 5AFI^30^). White contour represents area of the Empty system (Fig. 3) with Gibbs energy lower than 10 kJ/mol.

The CS width ranges from 0.4 to 1.2 nm regardless of the type of peptide accommodated in the tunnel. The global minimum of 0.33 nm was observed in sub-microsecond MD simulations of the empty ribosome, and roughly corresponds to the diameter of a water molecule. Because CS width is measured as a nucleus-to-nucleus distance, the true gap between the closest residues is smaller by the sum of the relevant van der Waals radii (C: 0.170 nm, N: 0.155 nm, O: 0.150 nm). Accounting for atomic size, we conclude that the CS can close completely, though this conformation is rare and accounts for only a few percent of simulation time.

Nevertheless, the CS spends a non-negligible fraction of time in conformations that would not permit NP passage. As a lower bound on NP diameter, we consider a polyglycine chain: a backbone alternating between planar peptide bonds and CH_2_ groups with two freely rotating bonds. The diameter of the narrowest cylinder capable of accommodating a glycine pentamer in the extended conformation (Φ = *−*120^*°*^, Ψ = 120^*°*^), including van der Waals radii, is approximately 0.67 nm. For typical side chains, the effective NP diameter is approximately 1.0 nm. As shown in Fig. 4C, the CS in the empty ribosome is narrower than 1.0 nm for approximately 75% of simulation time, implying that a significant fraction of conformations would sterically impede passage of a typical NP.

The CDF also informs the upper end of the CS-width distribution. The diameter of the smallest enclosing cylinder for an idealized *α*-helix is 0.88 nm, and 1.11 nm for polyglycine and polyalanine helices, respectively. Our simulations suggest that translocation of the simplest helical NPs would be geometrically compatible with the CS: the empty-tunnel CS exceeds 1.0 nm for approximately 25% of simulation time, sufficient to accommodate a narrow helix. However, *α*-helices bearing bulkier side chains would require local unfolding prior to CS passage, according to our model. In our MD simulations of Gly11 and Ala11 systems, no *α*-helix formation was observed within the simulation time. Short *α*-helices near the PTC have been reported in several stalling peptides, including VemP ^39,58^ in bacteria and hCMV^59^ in eukaryotes. However, none of the known *α*-helical NP fragments at the PTC belongs to the N-terminal portion of the protein; in all documented cases, the helix is formed by residues synthesized late in the sequence, meaning the N-terminus has long since passed the CS before the helix forms. The CDFs therefore imply that helix formation near the PTC does not necessarily preclude CS passage.

### 3.3 Peptide dynamics in the pre-constriction region

We simulated ribosomes with varying tunnel occupancy, using two model peptide systems with the simplest side chains to characterize NP dynamics in the pre-CS region. We expect that more realistic N-terminal sequences would exhibit slower dynamics, modulated by the chemical properties and folding propensities of their constituent residues.

To visualize the motion of tunnel contents and CS residues, we projected the positions of non-hydrogen atoms onto a cross-section of a cylindrical cutout aligned with the ribosomal tunnel axis (Fig. 2; see Methods). Projections for the empty tunnel and four peptide systems are shown in Fig. 6, where the contour lines enclose the regions occupied by atoms for 95% of simulation time.

**Figure 6:**
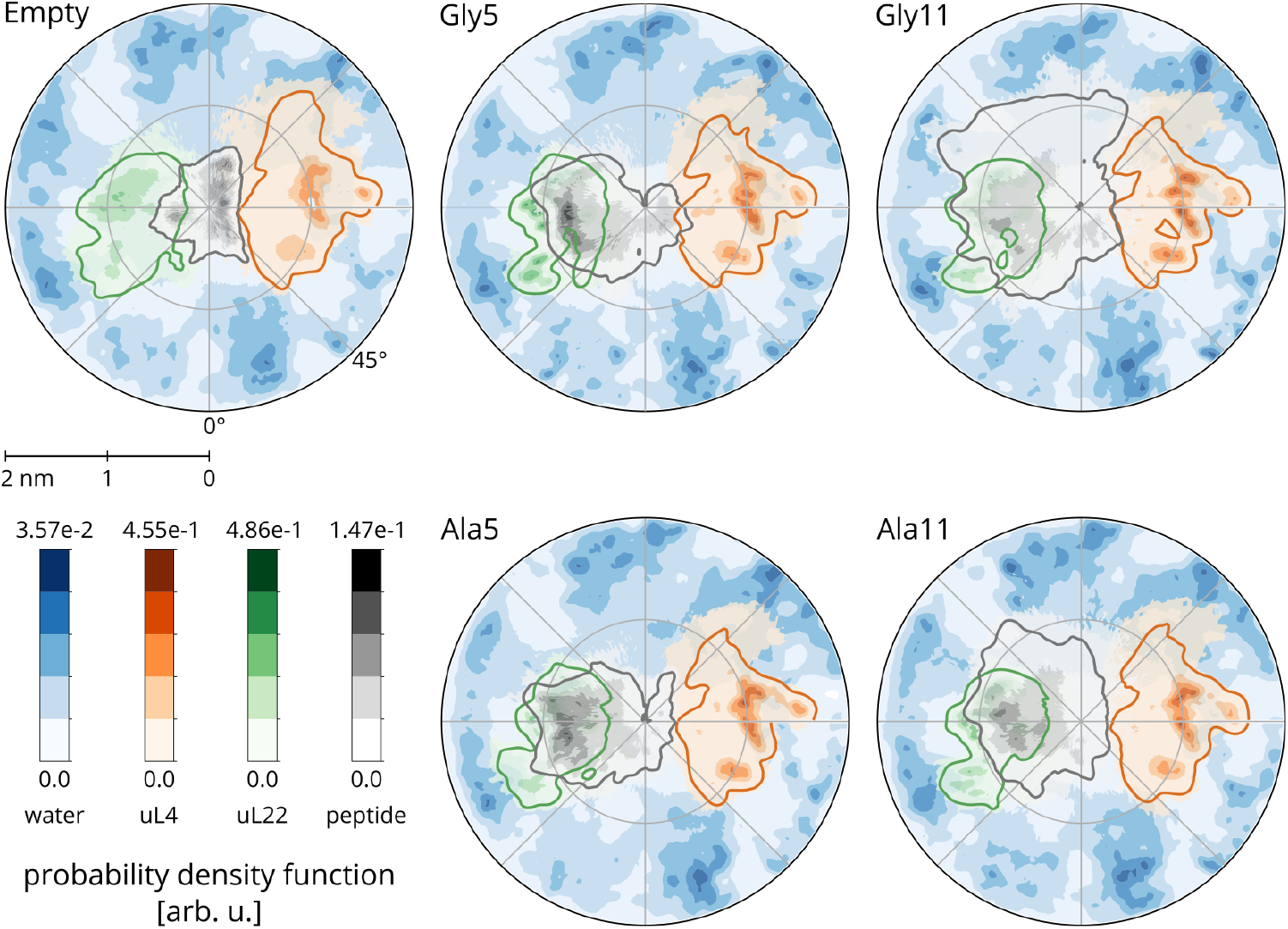
Probability density functions of non-hydrogen atoms’ positions within a cylinder (Fig. 2). The lines enclose 95% of the density.

A striking feature of these projections is a consistent preference of the NP to form an encounter complex with uL22, irrespective of NP length or composition. The probability density functions (PDFs) of NP positions show a clear maximum near uL22, indicating that uL22 is the primary binding site of the N-terminal formylmethionine. Because uL22 is evolutionarily younger than uL4, this preferential interaction may have acquired functional importance at a later stage of ribosome evolution. The potential role of the formyl group in driving this selectivity warrants further investigation.

NP flexibility follows intuitive trends: longer NPs are more flexible than shorter ones, and polyglycines are more flexible than polyalanines. This is reflected in the gray regions of Fig. 6, which grow with increasing NP length, and in the RMSF profiles (Fig. 7A). Additionally, for Gly11 but not Ala11, we observe folded-back conformations in which the N-terminal residue turns toward the PTC, as evidenced by short end-to-end distances (Fig. 7B). Similar behavior was observed in our previous study of peptide dynamics in carbon nanotubes, ^60^ where peptides were able to reverse direction despite strong confinement.

**Figure 7:**
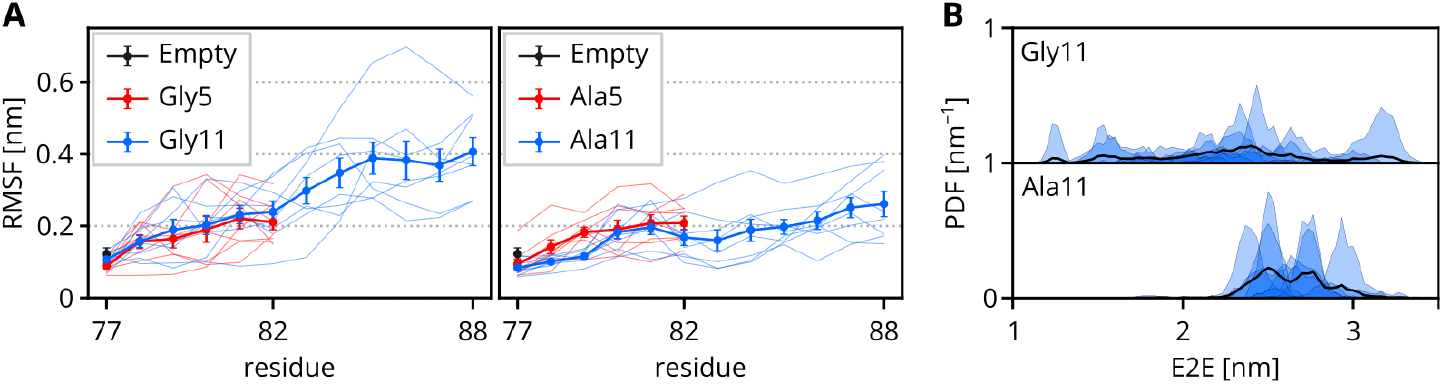
A) Root-mean-square fluctuations (RMSFs) of nascent polypeptide C_*α*_ atoms. Thin lines represent individual trajectories; thick lines are mean values with standard errors of the mean shown as error bars. B) Probability density functions (PDFs) of the end-to-end (E2E) Euclidean distance between the terminal residues of the nascent polypeptide. Blue areas represent individual trajectories; thick black lines are PDFs of all available data.

The folded-back conformations sampled by Gly11, in which the N-terminal residue approaches the PTC, raise the question of whether such configurations have functional consequences. One possibility is that a fully reversed NP could transiently occlude the PTC or interfere with the accommodation of incoming aminoacyl-tRNA, introducing a stochastic delay in elongation. This would be consistent with the translational ramp hypothesis, which posits that slow translation of early codons serves to prevent ribosome collisions and allow co-translational folding of downstream domains. ^61^ Indeed, Verma *et al*. observed^62^ a strong dependence of the protein yield on the codons 3–5. The difference observed here for Gly11 and Ala11 in our simulations is in line with the conclusion that the ramp is controlled at the NP level, more precisely by the conformational dynamics of short NPs.

Water density profiles were comparatively insensitive to tunnel occupancy. While water density was locally reduced in regions occupied by the peptide, the broader water distribution patterns remained largely unchanged across systems (Fig. 6).

The presence of an NP in the tunnel shifted the CS toward wider conformations relative to the empty ribosome, as seen in the Gibbs free-energy profiles (Fig. 5) and corresponding CDFs (Fig. 4C). This shift amounts to approximately 0.2 nm and is already apparent for the shorter peptides Gly5 and Ala5, suggesting that the CS can accommodate the NP passage by adaptively widening. This potentially allows larger NPs to traverse the tunnel without fully unfolding.

## 4 Conclusions

The ribosomal exit tunnel has long been depicted as a rigid conduit connecting the PTC to the ribosome surface.^63,64^ This view has been shaped largely by the nature of the experimental methods used to study ribosome structure: both X-ray crystallography and cryo-EM yield time- and ensemble-averaged models that suppress information about the dynamics. While these techniques have been indispensable for establishing the overall architecture of the tunnel, each deposited structure represents a single snapshot of what is, in reality, a continuously fluctuating molecular environment. Our work provides a quantitative picture of the conformational dynamics, focusing on the CS, the narrowest segment of the tunnel.

By analyzing 222 experimental structures of the *E. coli* ribosome, we showed that the CS exhibits substantial conformational variability even across the ensemble of static models available in the PDB. This variability is dominated by the side chains of a small number of residues, most notably Arg92 of uL22, which emerges from our analysis as a molecular gatekeeper of the CS, a function of Arg92 proposed previously in an early MD-simulation study. ^65^ Arg92 is simultaneously the most flexible residue at the CS and the one most frequently forming the closest contact with the opposing uL4 tip. Its long, positively charged side chain is capable of sampling multiple rotameric states, effectively scanning the width of the tunnel on the sub-microsecond timescale. The fact that Arg92 is highly conserved in prokaryotic ribosomes, and fully conserved in eukaryotes ^11^ suggests that this gating function is not an accidental byproduct of the local amino-acid composition but rather a selected feature of the translational machinery.

Unbiased all-atom MD simulations of the complete *E. coli* ribosome confirmed and extended these observations. The CS sampled widths ranging from fully occluded to conformations wide enough to accommodate a narrow *α*-helix, all within 200 ns of simulation time. This dynamic range is remarkable and has direct implications for nascent polypeptide translocation. Even the simplest peptide, polyglycine in an extended conformation, requires a CS opening of at least 0.67 nm, a threshold that is not always met. These findings suggest that CS passage is not a barrierless process but rather one that requires transient coordination between the conformational state of the CS and the geometry of the incoming peptide segment.

An important finding is that the presence of a nascent polypeptide in the tunnel, even one as short as five residues, shifts the CS toward wider conformations by approximately 0.2 nm relative to the empty ribosome. This adaptive widening indicates that the tunnel responds to its contents, rather than acting as a rigid conduit. Such responsiveness could serve a regulatory function, ensuring that the CS does not permanently obstruct the tunnel once peptide synthesis has begun, while still presenting a barrier to the earliest stages of translocation. This observation is also consistent with reports that macrolide antibiotics can modulate the conformation of the uL22 extended loop. ^13^ Consequently, an underappreciated mechanism of macrolide action may be the modulation of CS dynamics, directly affecting the ability of the exit tunnel to translocate NPs.

The preferential association of the N-terminal formylmethionine with uL22, observed consistently across all systems regardless of length or composition, identifies uL22 as the primary initial interaction partner of the NP. Given that uL22 is evolutionarily younger than uL4,^10^ this interaction may represent a relatively recent functional acquisition in the evolution of the ribosome. Whether the formyl group itself contributes to this selectivity, or whether the preference is dictated by the local electrostatic and geometric environment, remains an open question for future studies.

Our observation that the 11-residue polyglycine chain can adopt folded-back conformations, in which the N-terminus reverses direction toward the PTC, is potentially significant for the early kinetics of translation. Such conformations could transiently interfere with tRNA accommodation or peptide bond formation, introducing stochastic pauses in elongation. Our simulations suggest that the conformational freedom of the shortest nascent peptides, governed by their amino-acid composition, may contribute to the translational ramp at the molecular level, complementing the codon-usage and mRNA-structure effects that have been the primary focus of ramp studies to date. The observation that polyalanine chains, with their slightly bulkier side chains, do not exhibit the same folding-back behavior highlights how even subtle chemical differences at the N-terminus can alter the early dynamics of translation.

The high conservation of key CS residues – Gly91 and Arg92 of uL22, Gly64 and Thr65 of uL4 – is well documented,^11^ but the conventional interpretation has focused on the preservation of a specific three-dimensional structure. Our results suggest a different perspective: what has been conserved is not merely the static geometry of the CS, but its dynamic range, the capacity to fluctuate between open and closed conformations on a biologically relevant timescale. The most conserved residues at the CS are not structurally rigid; Arg92 is the most flexible residue in the region, and Gly91, nearly invariant across bacteria, is best ratio-nalized by the backbone flexibility it confers rather than by any structural role. The pattern of conservation thus favors a tunable gate over a rigid aperture, consistent with the need to accommodate diverse NP sequences while retaining the capacity to transiently impede passage. Testing this hypothesis will require comparative analyses of CS dynamics across phylogenetically diverse ribosomes, combined with mutational studies that selectively alter side-chain length or flexibility at conserved positions.

## Acknowledgment

We thank Tereza Svatoňová, who tested simulations models during her diploma thesis, and members of the Kolář research group at UCT Prague for valuable discussions.

## Authors’ contribution

MC prepared and analyzed the PDB data set, and setup the initial simulation systems; HM updated the simulated systems, performed simulations, and analyzed them; MK designed and supervised the research, and acquired funding; all authors interpreted the results, and drafted and finalized the manuscript.

## Data and software availability

The list of PDB structures analyzed, input simulation data, output data and the analysis scripts used to generate the figures are available online at https://github.com/kolarlab/mcgrath-constriction.

